# Kernel weight and source/sink ratio determination of temperate maize hybrids with different end uses under contrasting environments

**DOI:** 10.1101/2024.09.06.611734

**Authors:** Yésica D. Chazarreta, Santiago Alvarez Prado, Maria E. Otegui

**Author notes:** Corresponding authors (Y.D. Chazarreta), (M.E. Otegui).

## Abstract

Maize (*Zea mays L.*) production in Argentina changed markedly during the last decade due to the widespread adoption of late sowings, expanding its productive area, and diversifying crop end-uses. This study evaluated environment (two years × two sowing dates) and management practices (two nitrogen levels) effects on kernel weight, its physiological determinants, source/sink ratios, and water-soluble carbohydrates in stem (WSCS) of eight temperate maize hybrids bred for different uses (3 graniferous, 2 dual-purpose, 2 silage). Crop growth simulations allowed the estimation of percent variation in WSCS remobilization (null, partial, or total) for different production systems (18 scenarios) and climate conditions (41 growing seasons). Nitrogen fertilization increased kernel weight in early sowings, with minimal effects in late sowings. WSCS remobilization during kernel filling was higher in late than in early sowings, with no differences among hybrid types. Regarding hybrid types, dual-purpose and silage hybrids showed the highest and the lowest kernel weight respectively, and graniferous hybrids had the highest source/sink ratio during the effective kernel-filling period. Simulations underscored the importance of sowing date and nitrogen supply on WSCS for irrigated and dryland maize farming systems in a temperate environment, with important implications for grain and silage production at the farm level.

## 1. INTRODUCTION

Maize (*Zea mays* L.) grain yield is a complex trait that can be defined as a function of the number of harvested kernels and their individual weight. Although kernel number is usually the component that explains most of the yield variations (Andrade et al., 1996; Otegui, 1995), both components may affect the final yield (Gambín and Borrás, 2010; Cerrudo et al., 2013). The environment strongly influences kernel number (D’Andrea et al., 2013; Ruiz et al., 2019), while kernel weight is a highly heritable trait (Reddy and Daynard, 1983, Sadras, 2007), although it varies markedly among species (Gambín and Borrás, 2010), genotypic groups within species (Alvarez Prado et al., 2013; Hisse et al., 2019) and genotypes within a genotypic group (Borrás et al., 2009; Gambín et al., 2007).

Dry matter accumulation in the kernels begins after ovary fertilization and is usually divided into three phases: the lag phase, the effective kernel-filling period, and the maturity stage (Bewley and Black, 1985; Fig. 1b). The lag phase is a relatively short period characterized by a rapid increase in the kernel water content with negligible dry matter accumulation (Westgate and Boyer, 1986). It is a period with active cell division, where the number of endospermatic cells and amyloplast involved in the starch deposition is defined (Reddy and Daynard, 1983; Jones et al., 1996). The effective kernel-filling period is characterized by a rapid dry matter accumulation and can be described by two components: the kernel-filling rate and the duration of the phase (Egli, 1998). The kernel volume and its water content reach their maximum values during this phase (Borrás et al., 2003; Gambín et al., 2007) and then decline coordinately with the increase in biomass deposition (Fig. 1b). During the third and last phase, the rate of biomass accumulation decreases and the final kernel weight is established (Egli, 1998; Borrás and Otegui, 2001). At this time, commonly known as physiological maturity, kernels achieve their maximum weight and enter into a quiescent state, followed by a net water loss, which mainly depends on the prevailing environmental conditions (Chazarreta et al., 2023; Martinez-Feria et al., 2019; Schmidt and Hallauer, 1966).

**Figure 1.**
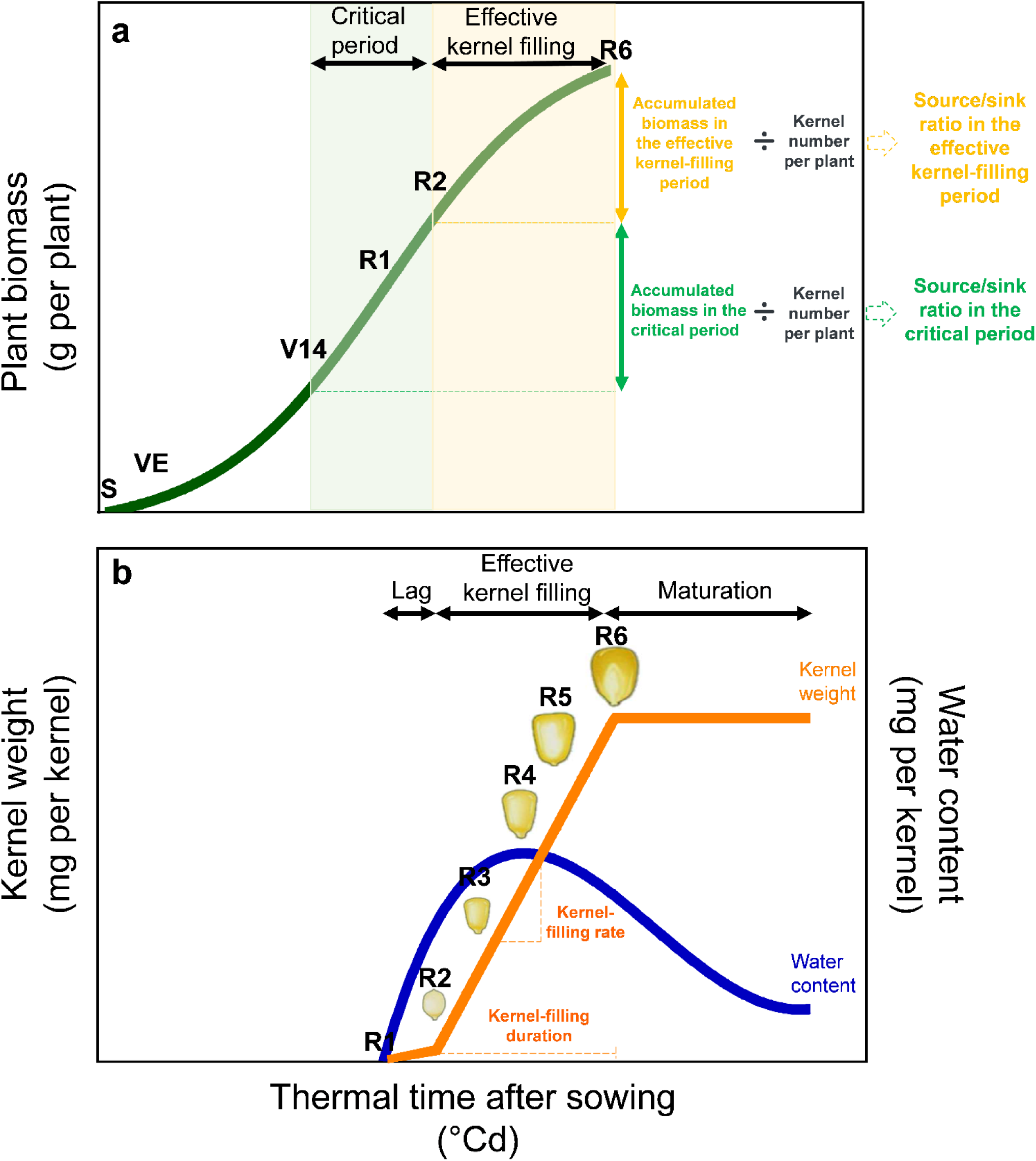
Scheme describing a) plant biomass accumulation and b) kernel dry matter accumulation (orange) and the kernel water content (blue) in response to the thermal time after sowing. In both figures, horizontal black arrows indicate the length of different phases. For biomass accumulation, the main physiological stages indicated along the cycle are sowing (S), emergence (VE), fourteen-leaf (V14), silking (R1), blister (R2), and physiological maturity (R6). For kernel dry matter accumulation, the indicated kernel stages are R1, blister (R2), milk (R3), dough (R4), dent (R5), and R6.

**Figure 2.**
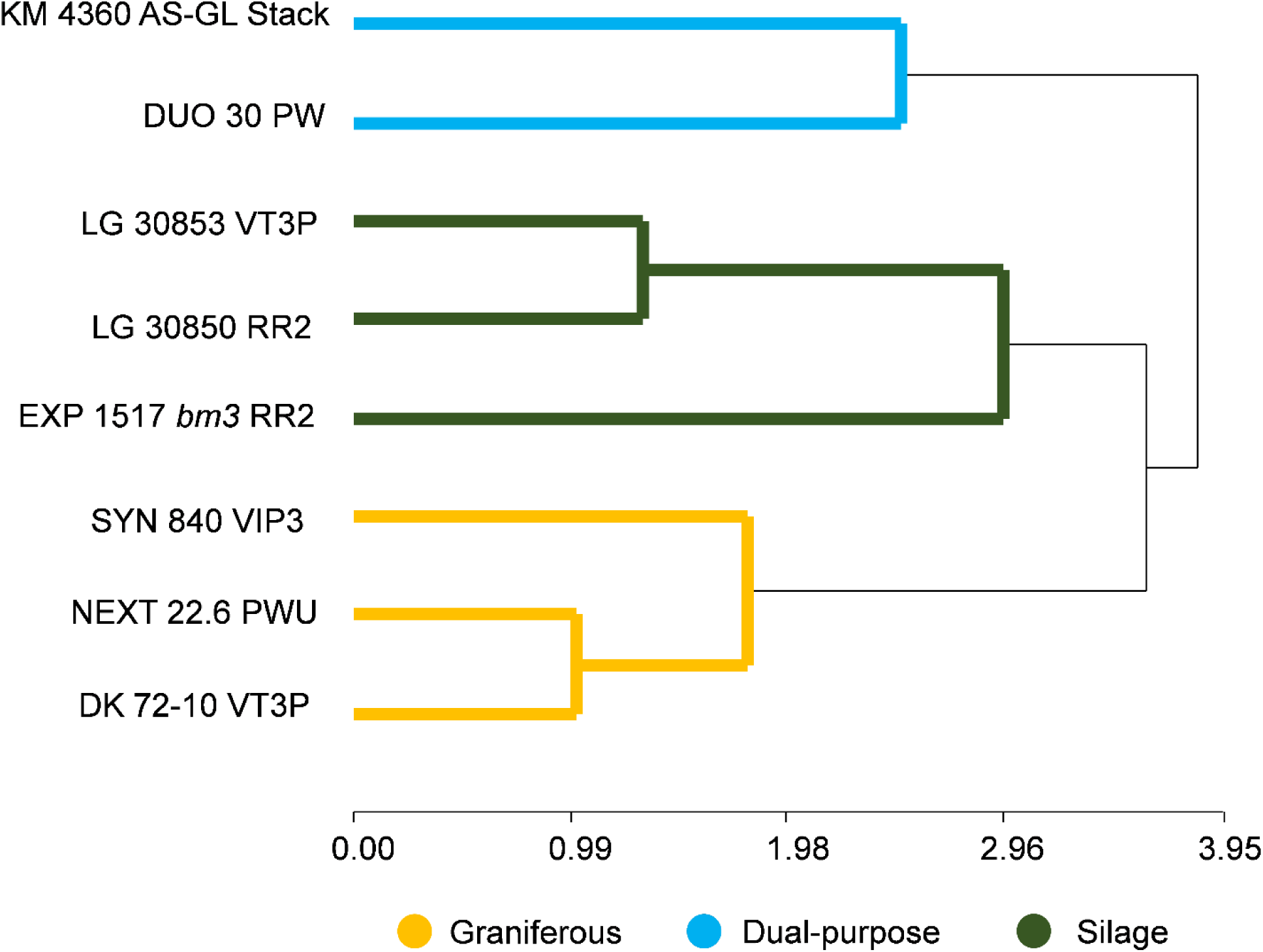
Cluster analysis for eight maize hybrids identifying three groups of hybrid types: graniferous (orange), dual-purpose (light blue), and silage (green).

Final kernel weight is affected by the environmental growing conditions explored during the effective kernel-filling period (Borrás and Otegui, 2001; Cirilo and Andrade, 1996). Assimilates for kernel filling are supplied by carbohydrates from current photosynthesis, nitrogen uptake from the soil, and from remobilization of reserves stored in vegetative organs (Gastal et al., 2012) in a ratio that depends on the genotype × environment combination. For early sown maize crops grown under non-limiting soil conditions, the main carbon assimilate source is photosynthesis with a marginal contribution of remobilization (Sekhon et al., 2016). Under such conditions, the final kernel weight did not increase much in response to increased assimilates availability per kernel (Borrás et al., 2004; Gambín et al., 2008). On the contrary, kernel growth is highly susceptible to source limitations during this phase (Borrás et al., 2004), which frequently reduce the final kernel weight due to a shortening of the grain-filling period (Olmedo Pico and Vyn, 2021; Badu-Apraku et al., 1983; Echarte et al., 2006). Nitrogen limitations during the crop cycle negatively impact kernel weight (Ruiz et al., 2022) through (i) reductions in the number of endosperm cells produced during the lag phase (Lemcoff and Loomis, 1994; Olmedo Pico et al., 2021), which can reduce the maximum kernel water content (Melchiori and Caviglia, 2008; Nasielski and Deen, 2019) and, consequently, the kernel-filling rate (Liu et al., 2011; Wei et al., 2019; Olmedo Pico and Vyn, 2021; Ren et al., 2021), or (ii) through a shortening of the kernel-filling duration (Olmedo Pico and Vyn, 2021). Under this situation, the contribution of reserves assimilates to the growing kernels markedly increases, mainly through remobilization of water-soluble carbohydrates stored in the stem (WSCS; Sadras et al., 2020) and N stored in the leaves and stems (Pan et al., 1986) during the vegetative stage. Under a low source/sink ratio, there is a high contribution of storage WSCS to the filling kernels (Uhart and Andrade, 1995), although this source of assimilates is insufficient to sustain the full kernel growth, which is interrupted when the reserves are depleted and there is no more contribution from direct photosynthesis (Rattalino-Edreira et al., 2014; Andrade and Ferreiro, 1996; Uhart and Andrade, 1995). Such conditions are frequently experienced by crops grown in environments characterized by a pronounced drop in temperature and irradiance during the kernel-filling period, like in high-latitude environments (Ruget, 1993; Kiniry and Otegui, 2000) and/or late sowing dates (Cirilo and Andrade, 1996; Bonelli et al., 2016).

Late sown maize has been associated with reductions in potential grain yield through decreases in the (i) number, size, and activity of the growing grains (Otegui et al., 1995; Cirilo and Andrade, 1994b, 1996), (ii) production of assimilates from photosynthesis during the kernel-filling period (Bonelli et al., 2016; Cirilo and Andrade, 1996; Tsimba et al., 2013), and (iii) duration and/or rate of kernel-filling (Chazarreta et al., 2021; Cirilo and Andrade, 1996). The effects of late planting over kernel weight and its physiological determinants turn particularly relevant for maize production in Argentina, where late-sown crops (i.e. near the summer solstice), as a single crop or double-cropped after a winter-grown species (e.g., pea, barley, or wheat as well as cover crops such as hairy vetch, ryegrass, or white oat), has increased from near zero in 2008-2009 to roughly 50% currently (Otegui et al., 2021). This trend promoted the expansion of the maize productive area toward the margins of the formerly main Central region (represented by the Rolling Pampas within the humid Pampas Region; Hall et al., 1992) and beyond (Otegui et al., 2021). The long distance from these regions to ports increases the transportation costs and consequently promotes on-farm maize usage, boosting alternative uses like silage, which currently represent ∼20% of the total cropped maize area in Argentina. Despite these trends, previous research has mostly focused on early sown maize for grain export (i.e., commodity; Otegui et al., 2021), and the interactive effects of the sowing date and nitrogen on the physiological determinants of the kernel weight and source/sink ratios have not been explored before. The main goal of this study was to assess the effect of the environment (year × sowing date) and management practices (nitrogen availability) on (i) kernel weight, (ii) its physiological determinants (kernel-filling rate and kernel-filling duration), and (iii) water-soluble carbohydrates in stem during the kernel-filling period for maize hybrids marketed for grain, dual-purpose (i.e. hybrids with good performance for both grain and silage purposes) or silage end-uses when grown under temperate conditions in Argentina. We hypothesized the explored environment (determined by the combination of years and sowing dates) and the management practices (N levels) will differentially affect the physiological determinants of kernel filling and the WSCS content depending on the hybrid type considered. For kernel weight, we expect delayed sowing dates to produce responses similar to those observed in high-latitude environments: reductions in the kernel weight due to decreases in the kernel-filling rate, a shortened kernel-filling duration, or both. Additionally, we anticipate lower kernel weight under the non-N-fertilized conditions, due to reductions in the kernel-filling rate and/or the kernel-filling duration. Different hybrid types are expected to exhibit differences in the physiological determinants of kernel filling, leading to an environment × hybrid type interaction for the kernel weight. Concerning WSCS, we expect that plants under late sowings will have lower values of WSCS near physiological maturity than the ones under early sowings.

## 2. MATERIALS AND METHODS

### 2.1. Field experiments

Field experiments were conducted at the Pergamino Experimental Station (33° 57’S, 60° 34’W) of the National Institute of Agricultural Technology (INTA), Argentina, on a silty clay loam soil (Typic Argiudol; USDA, 2014). A combination of two growing seasons (2019/2020 and 2020/2021, hereafter 2019 and 2020, respectively) and two sowing dates within each growing season (22-Oct and 19-Dec in 2019 and on 16-Oct and 18-Dec in 2020) were tested. Experiments planted in October were considered early, and those planted in December were considered late sowing date environments. Each combination of growing season × sowing date was considered an environment (hereafter 2019 Early, 2019 Late, 2020 Early, and 2020 Late). Genetic material consisted of eight F1, genetically modified (i.e., GMO) maize hybrids currently commercialized for grain (DK 72-10 VT3P, SYN 840 VIP3 and NEXT-22.6 PWU), silage (EXP 1517 *bm3* RR2, KM 4360 AS-GL Stack, LG 30850 RR2, LG 30853 VT3P) or dual-purpose (DUO30 PW). Hybrids were selected among those most representative of genotypes presently sown in the region for each purpose. The only exception was EXP 1517 *bm3* RR2, a pre-commercial hybrid included because of its brown midrib characteristic that was not present among the rest. At each environment, treatments consisted of the combination of hybrids and two nitrogen (N) levels: (i) no N added (N0), and (ii) full N, fertilized with 250 kg N ha^-1^ (N250) applied as urea at the fifth-leaf stage (V5; Ritchie and Hanway, 1986). Inorganic N levels (N-N0_3_) in the topmost 60 cm soil profile at sowing were 97 (Early 2019), 142 (2019 Late), 55 (2020 Early), and 80 kg N ha^-1^ (2020 Late). Organic matter levels at the experimental site were 3.1 and 3.4 % for 2019 and 2020 respectively.

At each environment, treatments were distributed in a split-split-plot design with three replications, with the sowing date in the main plot, N levels in the sub-plots, and hybrids in the sub-sub-plots (herein termed plots). Each plot consisted of six rows separated at 0.52 m and 7 m in length. Plots were hand-planted at a rate of 3-4 seeds per site and thinned in V2-V3 to obtain a single stand density of 9 plants m^-2^. Severe water stress was prevented through supplementary sprinkler irrigation. Crops were kept free of weeds, pests, and diseases using recommended chemical controls in all the experiments.

### 2.2. Measurements and calculations

Weather data were obtained from an automatic station located at less than 1 km from the experimental site and included daily values of mean air temperature (°C), mean solar incident radiation (MJ m^-2^ d^-1^), and rainfall (mm). Solar radiation was converted to photosynthetically active radiation (PAR) by multiplying by 0.45 (Monteith, 1965). Daily thermal time (in °Cday) was calculated as the difference between the daily mean air temperature (°C) and a base temperature (T_b_). The T_b_ was 8°C before silking (Ritchie and Nesmith, 1991) and 0°C from silking onwards (Muchow, 1990). For thermal time calculations, maximum daily temperatures exceeding 34°C were considered as 34°C (Cicchino et al., 2010).

The emergence date was recorded for each plot when 50% of plants reached this stage (VE). Then, five consecutive plants were tagged in the central row of each plot at V3. Silking date (R1, at least one silk visible in the apical ear) of each tagged plant was recorded. Non-destructive allometric models (Borrás and Otegui, 2001; Vega et al., 2000) were used to estimate the shoot biomass per plant at V6, V14 (*ca.* R1 – 15 days), R1, and blister R2 stage (R1 + 15 days; Fig. 1a). Models included the following variables: (i) stem volume, calculated from plant height to the uppermost visible collar and mean stem diameter at the base of the stalk (average of maximum and minimum values) and (ii) maximum ear diameter (at stages R1 and R2). Non-ear biomass (i.e., stem + leaves) was estimated with the stem volume, and earshoot biomass at R1 and R2 was estimated with the maximum ear diameter. Tassels were included with non-ear biomass. To build the allometric models, destructive samplings of nine plants from each hybrid × nitrogen × environment combination were carried out at V6, V14, R1, and R2, and the same morphometric variables were measured on them (Maddonni and Otegui, 2004). These plants were individually dried in a forced-air oven at 60 °C until constant weight and weighed to obtain non-ear and ear biomass. Linear (plant) and exponential (ear) models were fitted between biomass and morphometric variables.

Tagged plants were individually harvested at physiological maturity (R6) and dried in a forced-air oven at 60 °C until constant weight to obtain the final shoot plant biomass (g plant^-1^) and the kernel number per plant (KNP). Accumulated shoot biomass during the critical period (V14-R2) was considered as the difference between total shoot biomass at R2 and V14 (Fig. 1a). Accumulated shoot biomass during the effective kernel-filling period was calculated as the difference between total shoot biomass at R6 and R2 (Fig. 1a).

The source/sink ratio was calculated for the critical period (SSR_V14-R2_) and the effective kernel-filling period (SSR_R2-R6_) from data corresponding to the tagged plants. SSR_V14-R2_ was obtained as the quotient between the increase in shoot biomass from V14 to R2 and KNP (Fig. 1a). SSR_R2-R6_ was obtained as the quotient between the increase in shoot biomass from R2 to R6 and KNP (Fig. 1a).

A total of 12-15 plants were tagged in each plot at their exact silking date (R1) and used to estimate kernel-filling progress. Beginning 15 days after silking and continuing up to commercial kernel moisture (i.e. 14.5%), the apical ear of one plant per plot was harvested every 7 (before physiological maturity) or 15 days (after physiological maturity). The whole earshoot (ear + surrounding husks) was immediately enclosed in a plastic bag and transported to the laboratory in an insulated container. Fifteen kernels were sampled from the central section of each ear and used for the immediate assessment of kernel fresh weight. Kernel dry weight (KW) was measured after drying the kernels in a forced air oven at 70°C for at least 7 days. Kernel-filling dynamic was described by a bilinear with plateau model fitted to KW evolution (Eqs. 1 and 2) using thermal time after silking as the time variable (Borrás and Otegui, 2001):

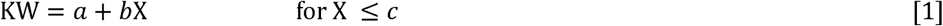

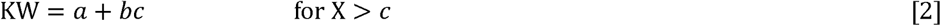

where *X* is the thermal time after silking, *a* is the intercept (in mg), *b* is the kernel-filling rate during the effective kernel-filling period (in °Cday^-1^) and *c* is the kernel-filling duration (in thermal time after silking). The maximum KW was the KW estimated by the *plateau* of the model, whereas the parameter *c* represented the physiological maturity date (in thermal time after silking; Fig. 1b). The duration of the lag phase (in °Cday) was estimated as the thermal time value where eq. [1] intercepted the x-axis (i.e. KW= 0). Kernel water content (KWC) was calculated by subtracting the kernel dry weight from the kernel fresh weight. Maximum kernel water content (MKWC) was determined for each environment × N × hybrid type combination by fitting a curvilinear model (Eq. [3]) as in Borrás et al. (2009):

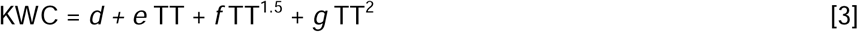

where KWC is water content and *d, e, f,* and *g* are model parameters.

Water-soluble carbohydrates in stem were measured every 15 days, beginning 15 days after silking (R2, blister stage) and up to R6. At each sampling date, one plant per plot was cut at ground level, and stem plus sheaths (hereafter termed stem) were separated from the rest of the plant. Plant height to the uppermost visible collar and diameter at the base of the stalk were measured on each sampled plant to estimate the stem volume. The stem of each plant was oven-dried until achieving constant weight, weighed, and ground. Water-soluble carbohydrates were measured in extracts from 0.1 g of dry tissue according to the methodology of Yemm and Willis (1954). The amount of water-soluble carbohydrates accumulated in the stem was calculated as the product of the stem dry weight and the concentration of water-soluble carbohydrates in the stem (Rattalino Edreira et al., 2014). The percentage of water-soluble carbohydrates accumulated in the stem (%) was calculated as the fraction of the stem weight corresponding to water-soluble carbohydrates. The remobilization of water-soluble carbohydrates in stem from R2 to R6 (in %) was calculated as follows:

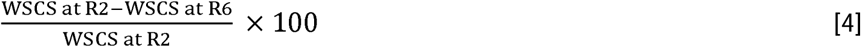

### 2.3. Statistical analyses and estimation of apparent stem reserves use in the predominant cropping-system scenarios

Cluster analysis (*k* = 3, Euclidean distance) was performed to classify the hybrids into their corresponding hybrid type: graniferous, dual-purpose, or silage. The analysis was based on their KNP, KW, kernel-filling rate, kernel-filling duration, SSR_V14-R2,_ and SSR_R2-R6_. The classification was performed using the hierarchical method based on the average linkage. Variables were standardized before conducting the cluster analysis. The joint interaction among the environment (growing season × sowing date), the N level, and the hybrid type was evaluated by principal components analysis (PCA). Biplots were constructed, where each environment × N × hybrid type combination was represented as a symbol and traits as vectors from the origin. The fixed effect of the environment, N, hybrid type, and their interactions were analyzed by performing analyses of variance (ANOVA), with block nested within the environment and hybrids nested within the hybrid type. A type-3 significance test for fixed effects was performed using α = 0.05, and their means were compared using Tukey adjustment. Cluster, principal components, and variance analyses were performed using Infostat (Di Rienzo et al., 2017). Regression analyses between variables and graphics were performed using GraphPad Prism version 9.0.0 (www.graphpad.com).

The effect of sowing date and initial soil conditions (i.e. plant available soil water (PASW) and N-NO_3_) on apparent stem reserves use (ASRU) during the effective kernel-filling period (ASRU_R2-R6_) was evaluated for the environment under study (temperate humid region) and the most frequent management practices used in this environment. These practices usually include the use of a single-cross maize hybrid of ca. 120 relative maturity under dryland farming, which can be sown early (September) or late (December) at a stand density of 75000 plants ha^-1^. In general, early sowings receive N fertilization and late sowings do not. For the evaluation, we used a modified version of MZCER980 (Mercau and Otegui, 2015; Otegui et al, 2021) included in the Decision Support System for Agrotechnology Transfer (DSSAT)-CERES-Maize V3.5 (Jones et al., 1998), and a climate series covering 41 growing seasons (1970-1971 to 2011-2012) obtained from the National Institute of Agricultural Technology (INTA; http://siga2.inta.gov.ar/<u>#/</u>) and the Prediction of Worldwide Energy Resource (NASA-POWER; https://power.larc.nasa.gov/data-access-viewer/). The ASRU_R2-R6_, in %, was estimated as in Eq. 4

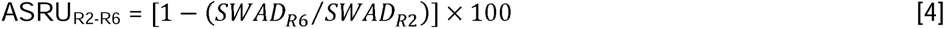

where SWADn in DSSAT represents aboveground stem dry weight at the indicated growth stages (R2 and R6).

Nine initial soil water × N scenarios were evaluated for each selected sowing date (15-Sep and 15-Dec). Water conditions included: (i) Irrigated, with the soil always at PASW > 90%, (ii) rainfed, starting with the whole profile (180 cm) at field capacity at sowing, and (iii) rainfed, starting with the whole profile at 50% PASW at sowing. For each water regime, three soil N conditions were evaluated: (i) no N restriction (Full N), and (ii) no N added during the cycle and initial soil N levels in the uppermost 45 cm as established by soil analyses in our experiments (i.e. 55 (N55) or 97 (N97) kg N ha^-1^ in early sowings and 80 (N80) or 142 (N142) kg N ha^-1^ in late sowings). For the rainfed conditions, automatic irrigation of 5 mm was added on sowing + 1 day to avoid excessive lengthening or failure of crop establishment.

## 3. RESULTS

### 3.1. Phenology and meteorological conditions

Early sowing dates reached silking (R1) between the end of December and the first week of January (31 Dec to 5 Jan in 2019 and 28 Dec to 3 Jan in 2020) while this stage was reached in the second half of February in late sowings (16 to 20 Feb in both growing seasons). Physiological maturity (R6) was reached during the second half of February and the first week of March in early sowings (16 to 24 Feb in 2019 and 23 Feb to 4 Mar in 2020), whereas this stage took place during the second half of April in late sowings (9 to 21 Apr in 2019 and 19 to 26 Apr in 2020). The duration of the post- silking period (from R1 to R6) was slightly longer for late (52 to 65 days) than for early (46 to 61 days) sowings. Details for silking and physiological maturity dates for each hybrid across environments are displayed in Table S1.

During the period that elapsed from VE to V14, mean air temperature was higher for late than for early sowings (7.5% and 15.6% for 2019 and 2020 respectively). By contrast, accumulated incident PAR during this period was higher for early than for late sowings (18.7% and 44.7% in 2019 and 2020 respectively; Table 1). During the critical period (V14 to R2), mean air temperature and accumulated incident PAR were similar between sowing dates (Table 1). Finally, during the effective kernel-filling period (R2-R6) mean air temperature was higher for early than for late sowing dates (14.1% and 17.9% for 2019 and 2020), as was accumulated PAR (20.5% and 34.0% in 2019 and 2020 respectively).

**Table 1.**
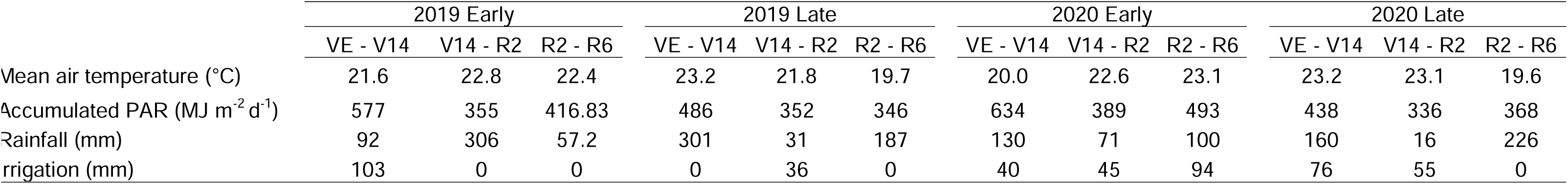
Mean air temperature, accumulated incident photosynthetically active radiation (IPAR), total rainfall, and total supplementary irrigation during three different periods for the four environments considered in the study.

The 2019 growing season belonged to the neutral phase of the ENSO (El Niño Southern Oscillation) phenomenon and 2020 to a moderate La Niña phase (https://ggweather.com/enso/oni.htm). Consequently, cumulative rainfall during the crop cycle (VE-R6) was higher in 2019 than in 2020 (51.2% and 29.1% for early and late sowing, respectively; Table 1).

### 3.2. Cluster analysis

The cluster analysis based on KNP, KW and its physiological determinants, and the source/sink ratios during the critical period and the effective kernel-filling period allowed for grouping the hybrids into three sets (Fig. 1). Hybrids DK 72-10 VT3P, NEXT-22.6 PWU, and SYN 840 VIP3 were clustered together as graniferous hybrids, whereas DUO 30 PW and KM 4360 AS-GL Stack could be considered as dual-purpose hybrids, and EXP 1517 *bm3* RR2, LG 30850 RR2, and LG 30853 VT3P as silage hybrids. This classification of the hybrids served as a framework for all subsequent analyses.

### 3.3. Kernel-filling and source/sink ratios

Kernel number per plant ranged from 416 to 574 across all environment × N × hybrid type combinations, and significant differences were detected on this trait for all main effects (p < 0.001; Table 2) as well as for the environment × N interaction (p < 0.01). The late-sowing environment caused a decrease in this trait during 2019 but not during 2020, and the graniferous hybrids presented a 12.0% greater KNP than dual- purpose and silage hybrids, which did not differ from each other (Table 2). On average, the N250 treatment increased KNP by 6.4%. However, the positive effect of N was significant only in the 2020 Early environment (+18.4% compared to the N0 one), without effect in the other environments.

**Table 2.**
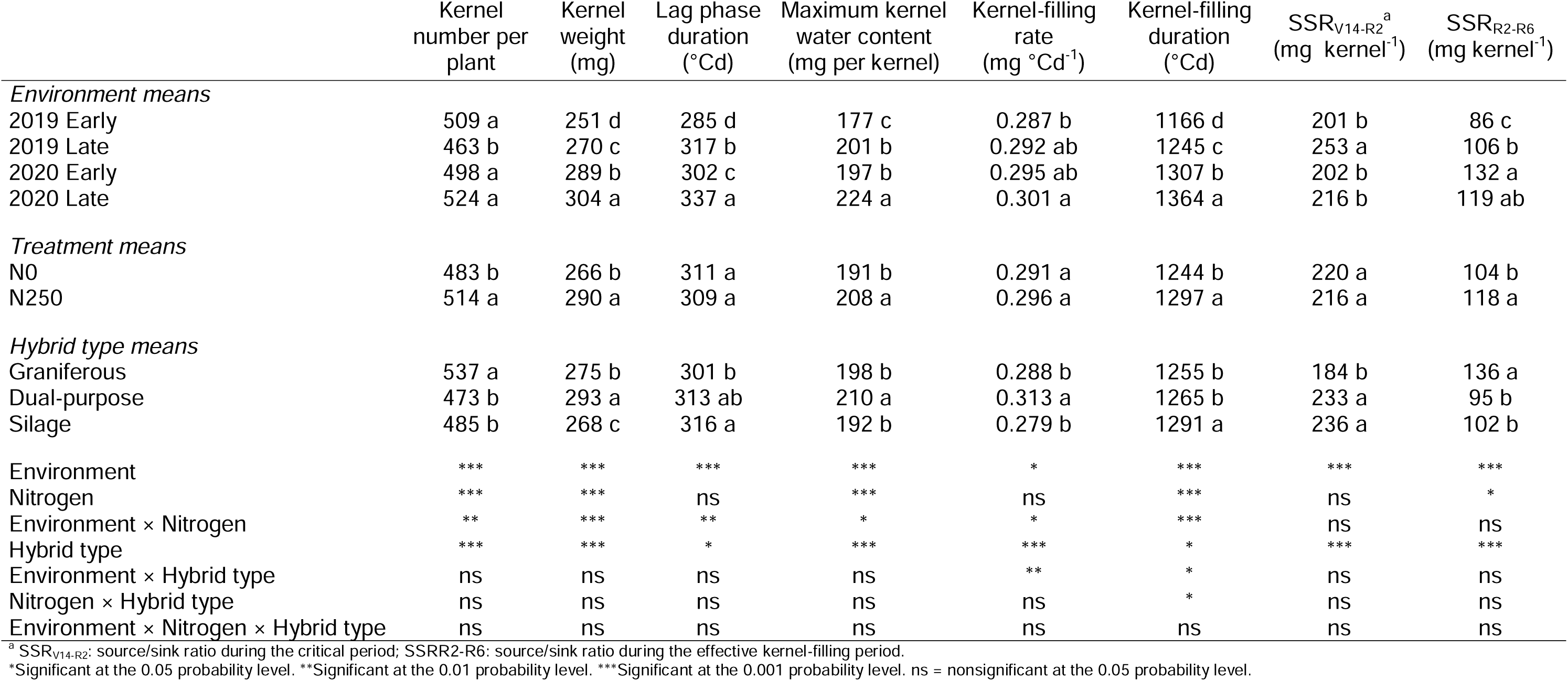
Means for the main effects of the environment (combination of years and sowing dates), the hybrid type (graniferous, dual-purpose, and silage), and the nitrogen level (N0, N250) on evaluated variables. ANOVA results are presented at the bottom of the table.

Significant main effects, as well as an environment × nitrogen effect (p < 0.001; Table 2), were also established for KW, which registered a maximum range of 215-327 mg. For this trait, however, all environments differed; the largest KW corresponded to the late sowing of 2020, and the smallest to the early sowing of 2019. Dual-purpose hybrids presented the highest KW, followed by the graniferous hybrids and finally by the silage ones. The N250 treatment significantly increased KW regarding the N0 one across all the environments, being the effect higher for early (17.1 and 13.0% for 2019 and 2020 respectively) than for late sowing date environments (3.3 and 4.0% for 2019 and 2020 respectively; Fig. 3).

**Figure 3.**
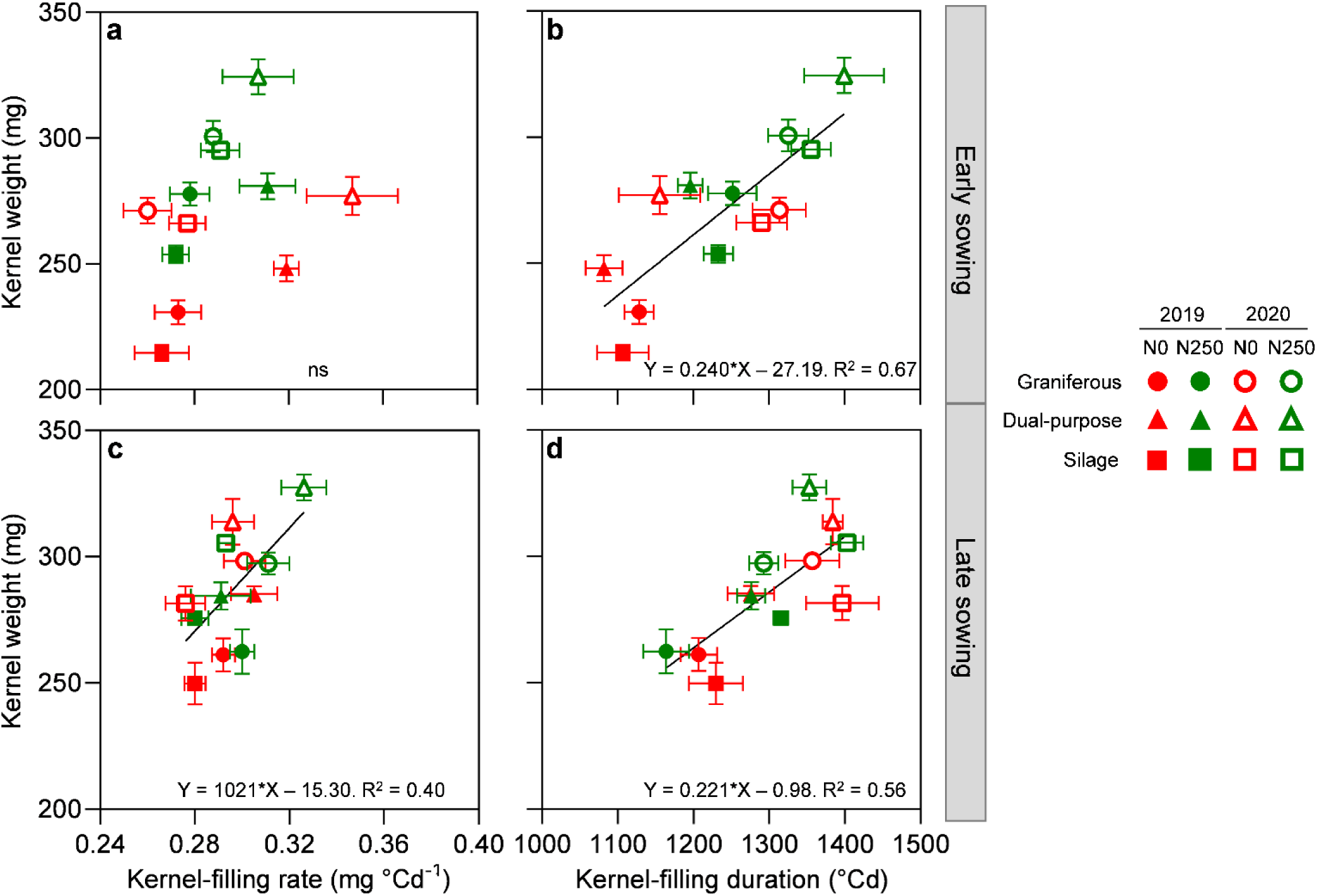
Relationship between kernel weight and its physiological determinants: (a, c) kernel-filling rate, and (b, d) kernel-filling duration. The upper panels correspond to early sowing dates (a, b), and the bottom panels to late sowing dates (c, d). Circles represent graniferous, triangles dual-purpose and squares silage hybrid types grown under different nitrogen levels (N0: no N added, in red; N250: 250 kg N ha^-1^, in green) during two growing seasons (2019: closed symbols; 2020: open symbols). Bars on each symbol indicate the standard error of the mean (SEM).

Regarding KW physiological determinants, the kernel-filling rate ranged from 0.260 to 0.347 mg °Cday^-1^. Significant environment (p < 0.05), hybrid type (p < 0.001) environment × nitrogen (p < 0.05), and environment × hybrid type (p < 0.01) effects were detected for this trait (Table 2). As for KW, the largest kernel-filling rate corresponded to the Late 2020 environment and the smallest to the Early 2019 environment, whereas the dual-purpose hybrid type showed the largest rate. The interaction effects detected that (i) the N250 treatment increased by 6.5% the kernel- filling rate concerning the N0 treatment only under the 2020 Late environment, with no effects on the rest of the environments (Table 2), and (ii) the dual-purpose hybrids exhibited a greater kernel-filling rate under early environments (15.8 and 17.2% in 2019 and 2020 respectively) than the graniferous and silage hybrids, which did not differ between them. The kernel-filling duration ranged from 1082 to 1403 °Cday^-1^ and varied across environments (p < 0.001), N levels (p < 0.001), and hybrid types (p < 0.05). Regarding the environments, the duration of this period was longer for the 2020 growing season than for the 2019 growing season, and late sowings had longer durations than the early ones (Table 2). The addition of N prolonged the kernel-filling period concerning the N0 conditions, and the silage hybrid had a longer kernel filling than the other types (Table 2). Significant environment × N (p < 0.001), environment × hybrid type (p < 0.01), and N × hybrid type (p < 0.05) effects were detected for this trait. The shortest duration was observed for the 2019 Early × N0 × dual-purpose hybrid type while the longest duration corresponded to the 2020 Late × N250 × silage hybrid type combination. Kernel-filling duration was longer under N250 than N0 conditions in early sowings (9.5 and 8.5% for 2019 and 2020 respectively), while N treatments did not affect the duration of the period in late sowings. Under the N250 treatment, silage hybrids showed a 5.4% longer kernel-filling duration than the graniferous ones, while dual-purpose hybrids presented an intermediate duration that did not differ from the other hybrid types. Hybrids did not differ in kernel-filling duration under N0 treatments. The increases in KW were linearly and positively related to (i) the increases in kernel- filling rate for late but not for early sowings (Figs. 3a and 3c), and (ii) the increases in kernel-filling duration for both early (Fig. 3b) and late (Fig. 3d) sowing date environments.

Duration of the lag phase ranged from 264 to 352 °Cday and differed among environments (p < 0.001), hybrid types (p < 0.05), and environment × N combinations (p < 0.01). On average, the lag-phase duration was longer in the 2020 than in the 2019 experiments, and for late than early sowings within each experiment (Table 2). Silage hybrids presented a slightly longer (4.9%) lag phase duration than the graniferous ones, while dual-purpose hybrids showed an intermediate lag phase duration that did not differ from the other hybrid types. The lag phase was 4.4% longer under the N250 than the N0 treatment in the 2020 Late environment, while N addition did not affect this trait in the rest of the environments.

The maximum kernel water content (MKWC) ranged from 160 to 240 mg kernel-1. As for the durations described above, the MKWC was higher in the second than in the first growing season, and in late than in early-sowing conditions (p < 0.001, Table 2). Dual-purpose hybrids exhibited a 7.6% greater maximum water content than the grain and silage hybrids, which did not differ for this variable (p < 0.001, Table 2). On average, the MKWC was 8.9% higher for the N250 than for the N0 condition (p < 0.001, Table 2), but the effect of N addition varied across the different environment × N combinations (p < 0.05). The MKWC was 18.3% greater for the N250 treatment in the 2019 Early environment, whereas this trait did not respond to N addition in the rest of the environments. Across environments, the increases in MKWC were linearly and positively related to the increases registered in the lag-phase duration (Fig. S1a), and the increases in kernel-filling rate were linearly and positively associated with the increases registered in the MKWC (Fig. S1b).

The source/sink ratio during the critical period ranged from 151 to 272 mg kernel^-1^ and differed among environments (p < 0.001) and hybrid types (p < 0.001). SSR_V14-R2_ was 25.8% greater in the 2019 Late than in the 2019 Early environment, with no differences between sowing dates in 2020. Silage and dual-purpose hybrids presented a 27.5% greater SSR_V14-R2_ than the graniferous ones, mainly explained by a lower KNP (Table 2). SSR_R2-R6_ ranged from 67 to 168 mg kernel^-1^ and differed among environments (p < 0.001), nitrogen treatments (p < 0.05), and hybrid types (p < 0.001). SSR_R2-R6_ was 23.0% greater in the 2019 Late than in the 2019 Early environment, with no differences between sowing dates in 2020. Averaged across environments, the N250 treatment increased by 12.9% the SSR_R2-R6_ respect to the N0 one. Graniferous hybrids presented a 37.7% higher SSR_R2-R6_ than the average of other types, which did not differ from each other. Lastly, kernel weight showed a significant (p < 0.05) and positive linear relationship with the SSR_R2-R6_ in early sowing dates (Fig. 4a). Considering its physiological determinants, this response to the SSR_R2-R6_ was not driven by the kernel-filling rate (p > 0.05; Fig 4b) but by the kernel-filling duration (p < 0.01; Fig 4c). Neither kernel weight nor its components were related to the SSR_R2-R6_ in late sowing environments (Fig 4 d, e, and f).

**Figure 4.**
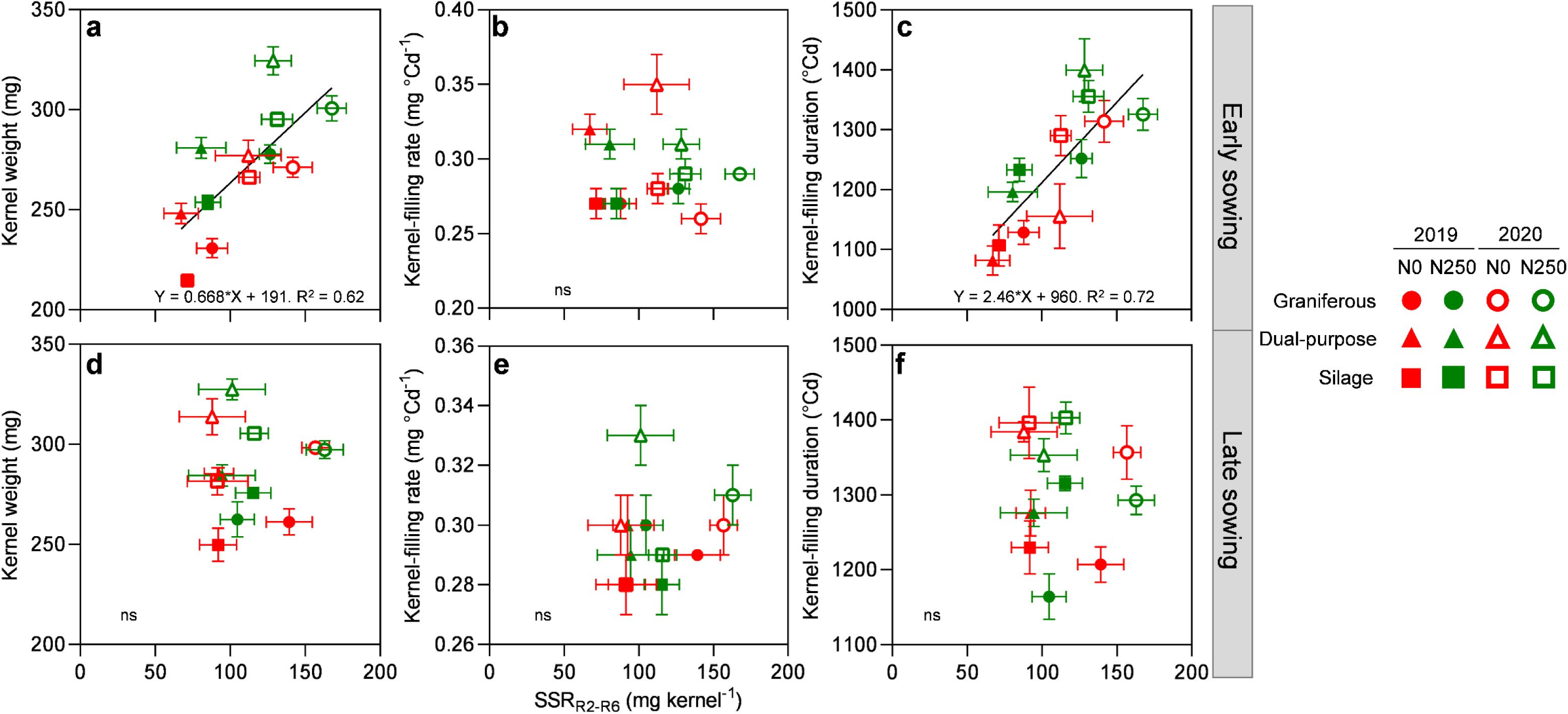
Response of kernel weight and its components, the kernel-filling rate, and kernel-filling duration, to the source/sink ratio during the effective kernel-filling period (SSR_R2-R6_). The upper panels correspond to early sowing dates (a, b, and c), and the bottom panels to late sowing dates (d, e, and f). Circles represent graniferous hybrids, triangles dual-purpose and squares silage hybrids grown under different nitrogen levels (N0: no N added, in red; N250: 250 kg N ha^-1^ in green) during two growing seasons (2019: closed symbols; 2020: open symbols). Bars on each symbol indicate the standard error of the mean (SEM).

### 3.2. Multi-traits associations

Principal component analysis was performed to evaluate the relationships among the attributes across the different treatments and environments. The analysis involved kernel number per plant, kernel weight, lag phase duration, maximum kernel water content, kernel-filling rate, kernel-filling duration, SSR_V14-R2,_ and SSR_R2-R6_. The first two components accounted for 72.0% of the observed variation, with 42.6% for the first PCA (PC1) and 30.4% for the second PCA (PC2; Fig. 5). Correlation coefficients between the variables and their associated probabilities are displayed in Tables S2 and S3, respectively.

**Figure 5.**
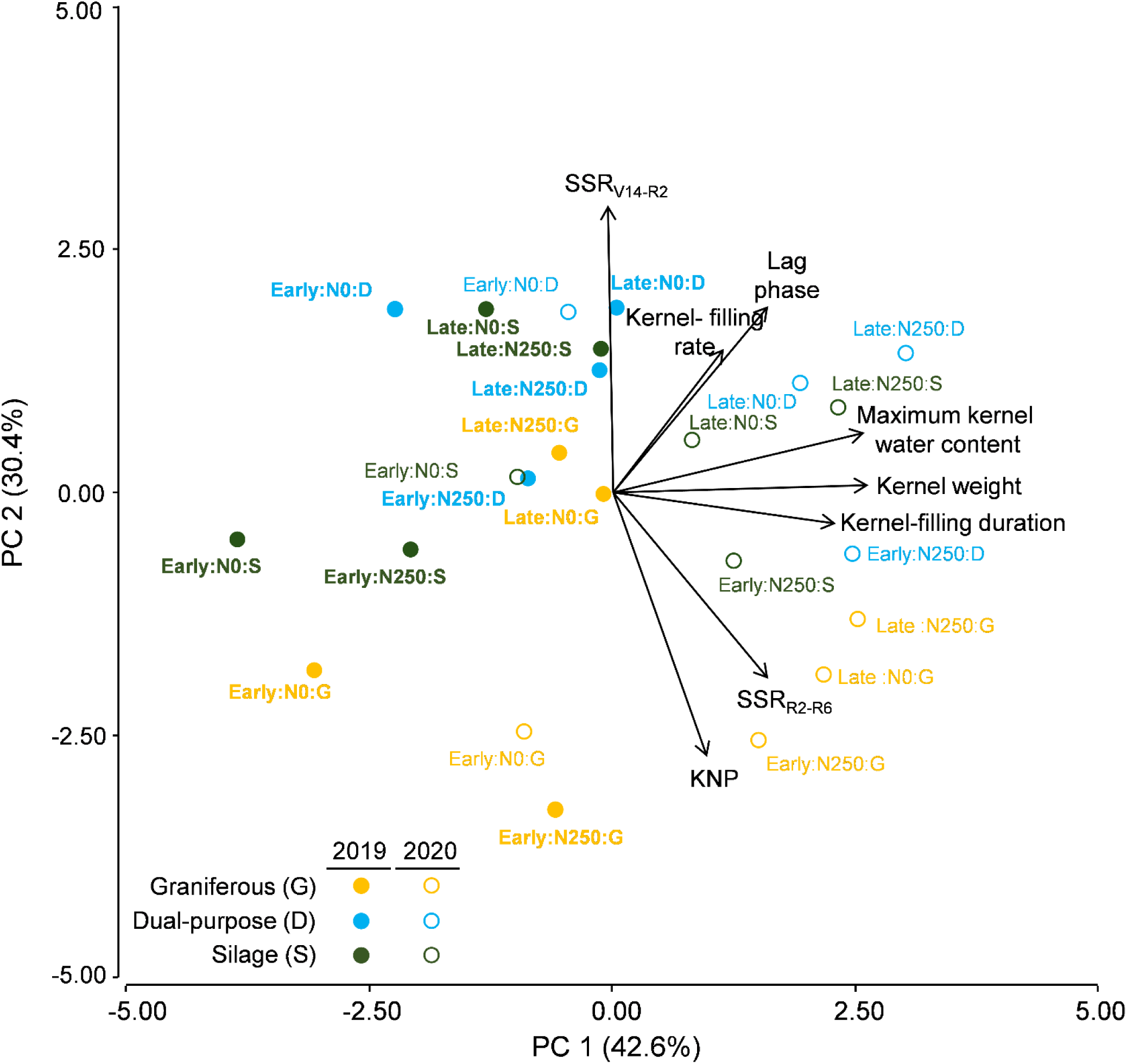
Biplot for the first two principal components (PC1 and PC2) of eight maize hybrids grouped into three types (graniferous in orange, dual-purpose in light blue and silage in green) grown under different combinations of year (2019: closed symbols with bolded labels; 2020: open symbols with regular labels), sowing date (Early and Late) and N level (N0: no N added; N250: 250 kg N ha^-1^). Traits in black solid vectors are for kernel number per plant (KNP), kernel weight, maximum kernel water content, lag phase and kernel-filling durations, source/sink relation during the V14-R2 period (SSR_V14-R2_) and source/sink relation during the R2-R6 period (SSR_R2-R6_).

The PC1 encompassed the variation observed in kernel weight, maximum kernel water content, and kernel-filling duration, with comparatively higher values of these variables towards positive values of the x-axis. The implicit projection of these variables in the opposite direction would indicate comparatively low values of them. The second principal component (PC2) encompassed the variation observed mainly in both SSRs, KNP, duration of the lag phase, and the kernel-filling period, with comparatively higher values of KNP and SSR_R2-R6_ towards negative y-axis values and comparatively higher values of the other three variables towards positive y-axis values (Fig. 5).

Kernel weight presented (i) a strong positive association (vectors in very acute angle; Fig. 5) with the maximum kernel water content (*r* = 0.89, p < 0.0001) and the kernel-filling duration (*r* = 0.82, p < 0.0001), and (ii) a slight and positive association (vectors in moderate acute angle; Fig. 5) with the duration of the lag phase (*r* = 0.41, p < 0.0001) and the SSR_R2-R6_ (*r* = 0.46, p < 0.05; Tables S2 and S3). The kernel-filling rate had a lower weight in the analysis because its vector was the shortest among all evaluated traits. Kernel number per plant was strongly and negatively associated with the SSR_V14-R2_ (*r* = -0.75, p < 0.0001; Tables S2 and S3) and positively associated with the SSR_R2-R6_ (*r* = 0.56, p < 0.01; Tables S2 and S3). On the one hand, the source sink/ratios during the V14-R2 and R2-R6 were negatively associated between them (*r* = -0.50, p < 0.05; Tables S2 and S3), a trend that was anticipated by the ANOVA analysis (Table 2). On the other hand, the correlation between grain yield components (i.e. KNP and KW) as well as between the physiological determinants of KW (i.e. kernel-filling rate and kernel-filling duration) was almost null (vectors at near 90°).

Among environments, most of the data of the 2020 growing season were displayed towards positive values of the x-axis (i.e. comparatively higher KW values) and those of the 2019 growing season were displayed in the opposite direction (i.e. comparatively lower KW values). Respect to hybrids, those of the graniferous type projected their comparatively higher records towards negative values of the y-axis, associated with higher values of KNP and SSR_R2-R6_, while dual-purpose and silage hybrids did it in the opposite direction in association with higher values of SSR_V14-R2_. Only data from 2020 combined high KNP with high KW values (5 cases located between the positive projections of both vectors). In most cases (4 out of 5) they corresponded to the N250 condition, particularly in the early sowing of all hybrid types (3 out of 4). The opposite condition (combination of low KNP and KW values) was dominated by 2019 data (4 out of 5), and silage (2 out of 5) or dual-purpose (2 out of 5) hybrids under N0 (3 out of 5), with no clear trend between sowing dates (the worst performance corresponded to non-fertilized dual-purpose hybrids sown early and silage hybrids sown late that year).

### 3.3. Water-soluble carbohydrates in stem dynamic

Beginning in R2 (*ca.* 15 days after silking), the percentage of water-soluble carbohydrates in stem ranged from 21 to 32% across all environment × N × hybrid type combinations and differed among environments (p < 0.001), N treatments (p < 0.05) and hybrids types (p < 0.05; Table 3). Early sowings started the effective kernel-filling period with higher percentages of WSCS than the late ones (+19.4 and +29.2% for 2019 and 2020, respectively; Table 3, Fig. 6). Plots under the N250 treatment started the effective kernel-filling period with a higher percentage of WSCS (7.2%) than the N0 ones (Table 3). Silage hybrids presented higher values of WSCS in R2 than the dual- purpose ones (6.5%), while grain hybrids presented intermediate values that did not differ from the other two hybrid types. At the R4 stage (*ca.* R1+30 days), the percentage of WSCS ranged from 17 to 28% and differed among environments (p < 0.01) and N treatments (p < 0.05; Table 3). Values of WSCS were 28.9% higher in early than in late sowing date environments in 2020, with no differences between sowing dates in 2019, and 9.7% higher for N250 than N0 treatments (Table 3, Fig. 6). Following the R5 stage (*ca.* R1+45 days), the percentage of WSCS ranged from 4 to 22% and significantly differed across environments (p < 0.001). Early sowings presented a significantly higher percentage of WSCS at this stage than the late sowings (76.9 and 40.3% for 2019 and 2020 respectively; Table 3, Fig. 6). Finally, the percentage of WSCS at R6 ranged from 4 to 21% and showed a significant environment × N (p < 0.05) interaction. The N0 treatment presented 52.5% higher WSCS than the N250 one in the 2019 Early environment (p < 0.05), with no differences between N treatments in the rest of the environments. Data of WSCS for R2, R4, R5, and R6 expressed in g per plant are displayed in Table S4.

**Figure 6.**
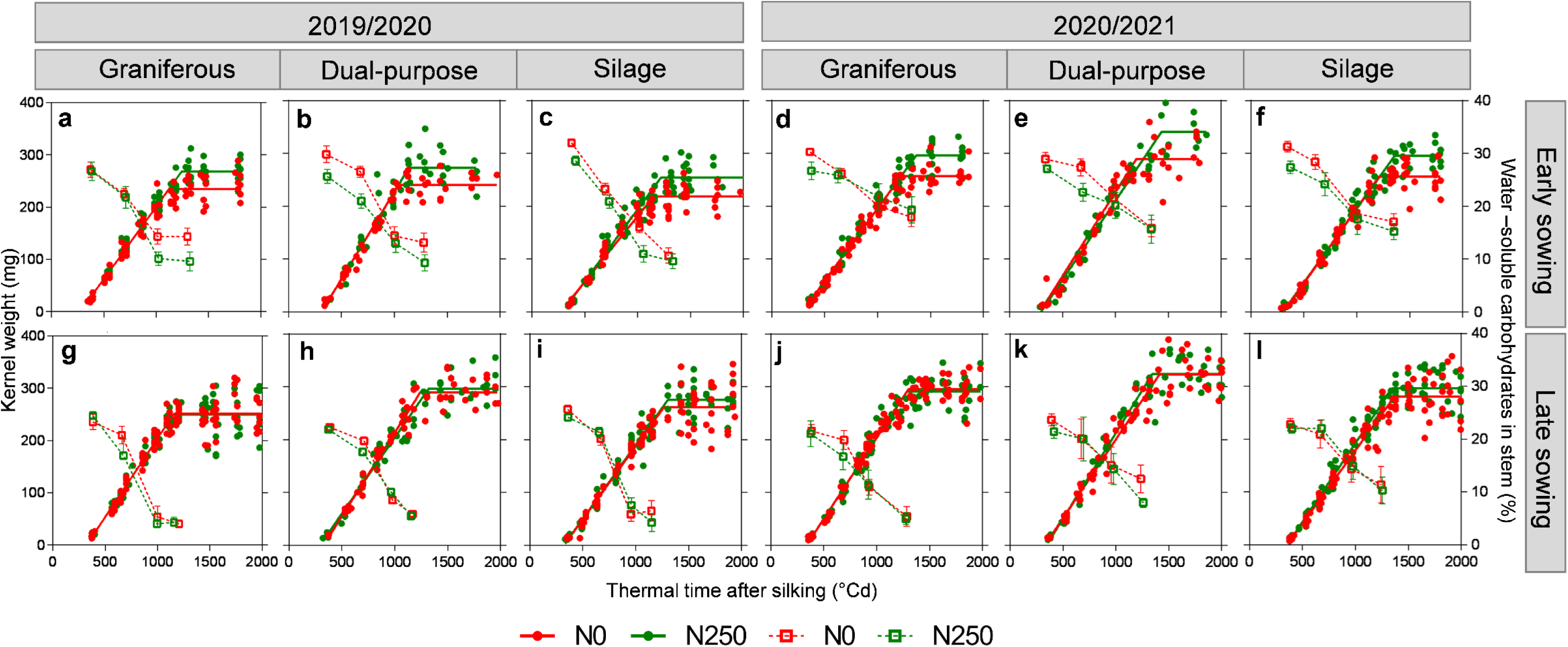
Evolution of the kernel weight (filled-circle symbols; left y-axis) and water-soluble carbohydrates in stem (open-square symbols; right y-axis) in function of the thermal time after silking for graniferous (a, d, g, j ), dual-purpose (b,e,h,k) and silage (c, f, I, l) maize hybrid types grown under different nitrogen levels (N0: no N added, in red; N250: 250 kg N ha^-1^ in green) during two growing seasons: 2019/2020 (a-c, g-i) and 2020/2021 (d-f, j-l). Upper panels (a-f) correspond to early sowings and bottom panels (g-l) to late sowings. For kernel weight, solid lines represent fitted models. For water-soluble carbohydrates in stem, dashed lines connect adjacent points and bars on each symbol indicate the standard error of the mean (SEM).

**Table 3.**
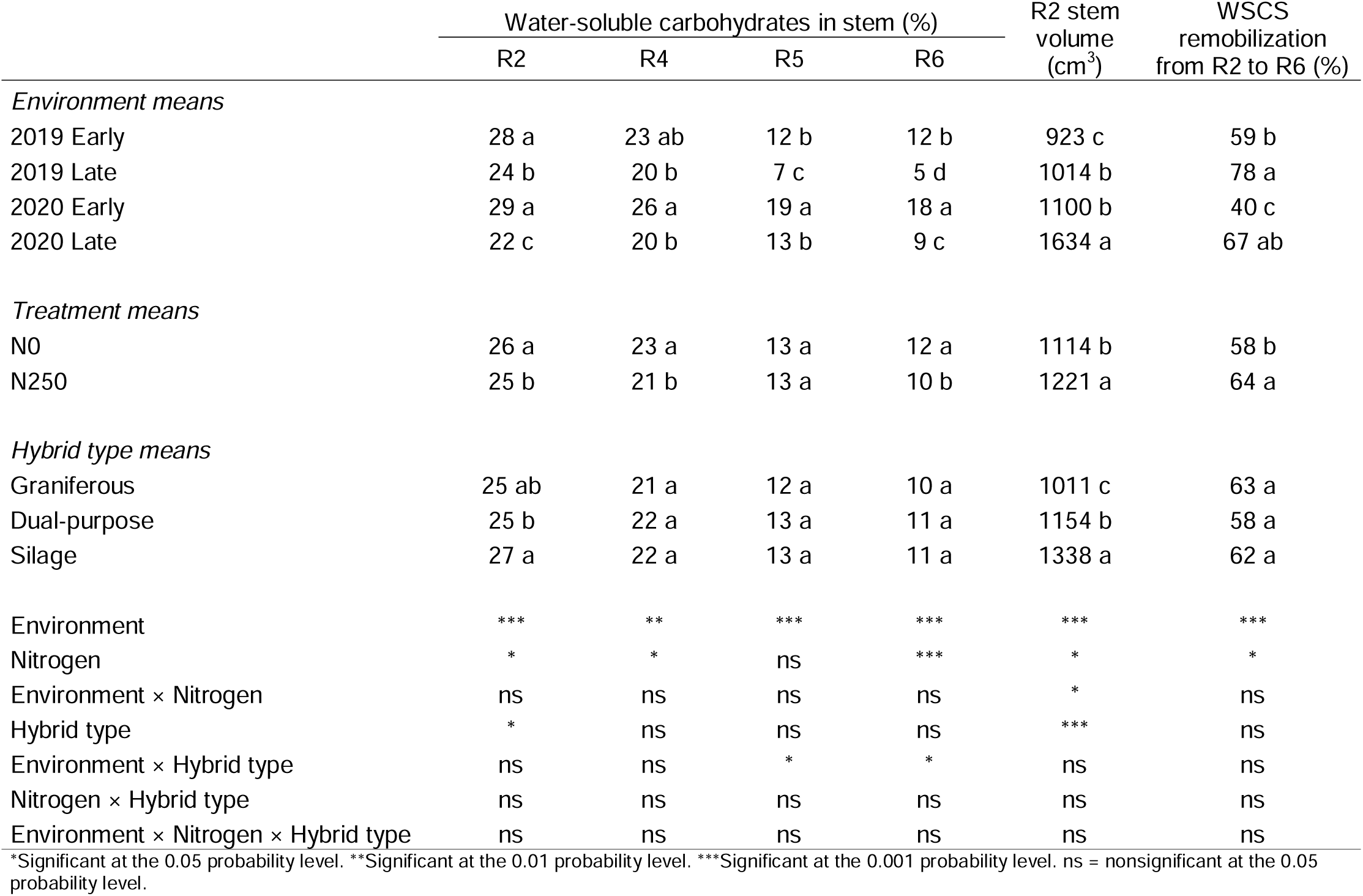
Means for the main effects of the environment (combination of years and sowing dates), the hybrid type (grain and silage) and the nitrogen level (N0, N250) of water-soluble carbohydrates in stem (WSCS) at R2 (*ca.* 15 days after silking; blister), R4 (ca. 30 days after silking; dough), R5 (ca. 45 days after silking; Dent), and R6 (ca. 60 days after silking; maturity), stem volume at R2, and remobilization of WSCS from R2 to R6. ANOVA results are presented at the bottom of the table.

Stem volume at R2 ranged from 810 to 1978 cm^3^, varying across environments (p < 0.001), N (p < 0.05), hybrid types (p < 0.05) and environment × N combinations (p < 0.05). Stem volume at R2 was largest for the 2020 Late environment and smallest for the 2019 Early environment, with the other two environments exhibiting intermediate values (Table 3). Average across growing conditions, the N250 produced larger stem volumes than the N0 one (+8.9%), and the order among hybrids was silage>dual-purpose>graniferous type (+15.9% for the first comparison and +14.1% for the second). The environment × N significant interaction detected that stem volume was 17.9% higher under the N250 than under the N0 treatment in the 2020 Late environment, with no differences between N treatments for the rest of the environments.

The WSCS greatly diminished during the effective-kernel filling period (Table 3; Fig. 6). The remobilization percentage of WSCS from R2 to R6 ranged from 30 to 87% and varied across environments (p < 0.001) and N treatments (p < 0.05). Late sowing date environments remobilized a greater percentage of WSCS from R2 to R6 than the early ones (32.1 and 67.8% in 2019 and 2020 respectively), while the WSCS remobilization from R2 to R6 was 8.9% greater under the N250 condition than under the N0 one (Table 3).

### 3.4. Estimation of stem reserves use for different crop management practices across years

The simulation of 18 different cropping-system scenarios across 41 growing seasons showed substantial variation in apparent stem reserves use during the active kernel-filling period (ASRU_R2-R6_; Fig. 7). The extreme situations concerning either null reserve use or completely depleted WSCS at R6 corresponded to the simulated early sowing of mid-September under no water deficit. On the one hand under potential conditions (i.e. with no water or N restrictions) and on the other hand under low initial soil N content (N55). For the former, simulations indicated that no ASRU_R2-R6_ could be expected in 42% of the growing seasons, whereas for the latter this value dropped to only 3% of the growing seasons. Additionally, cases with stem biomass loss larger than the estimated levels of WSCS at R2 (i.e. complete depletion of WSCS during active kernel filling) were almost null for the former (only 2%) and largest for the latter (51%). The situation was similar under rainfed conditions. Firstly, because the proportion of cases with null ASRU_R2-R6_ was highest (39%) and lowest (12%) under early sowing dates and for crops that started their cycle with soils at field capacity. The former corresponded to crops that experienced no N limitation along the cycle and the latter to those with low initial soil N conditions (N55) that were not fertilized with N. Secondly, because the proportion of cases with complete WSCS depletion was minimum (2%) and maximum (44%) for the same early sowing date scenarios (i.e. those that started with high initial PASW), the former without N restrictions and the latter with low initial soil N and no added N.

**Figure 7.**
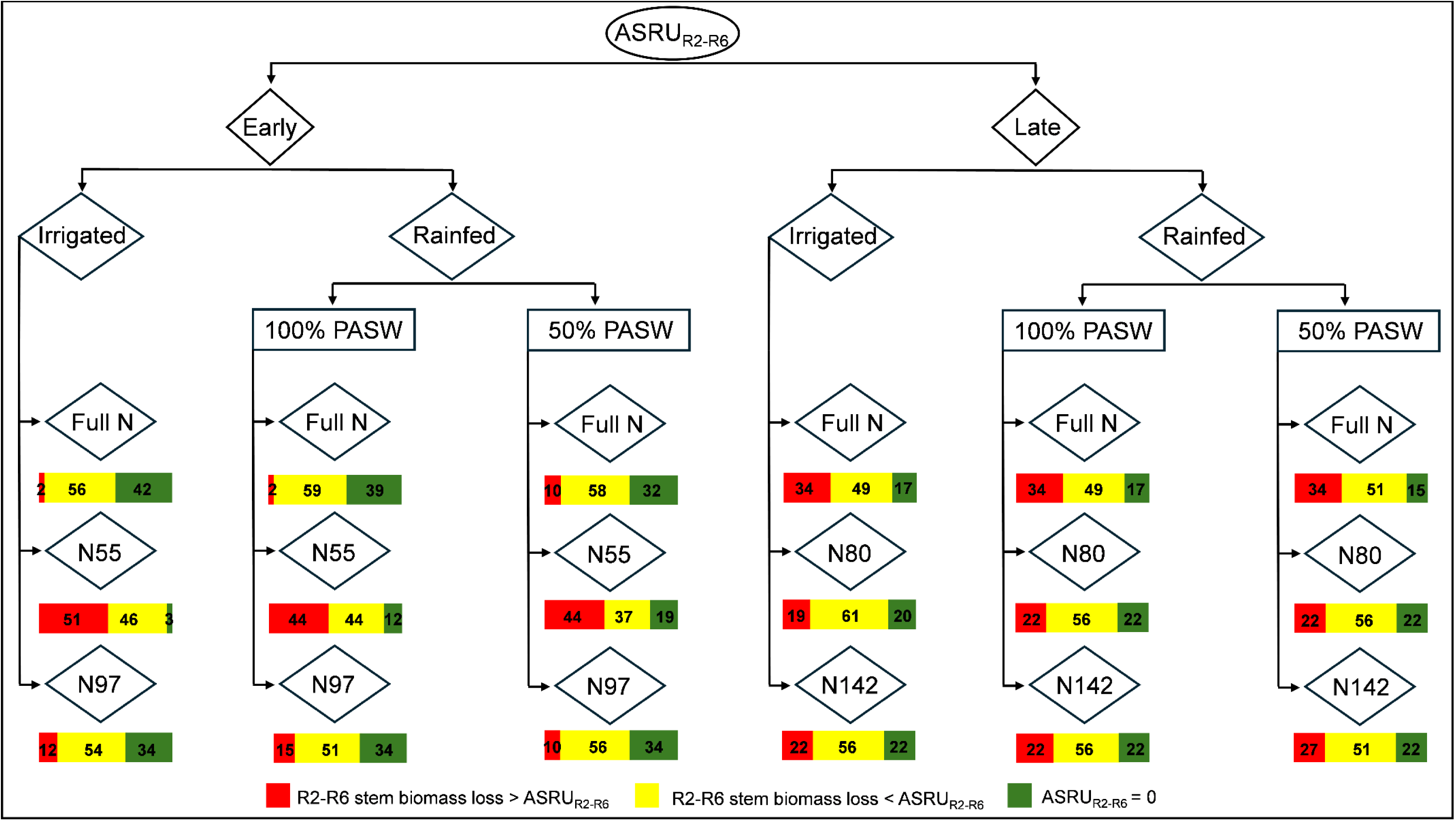
Flow chart illustrating the trends in apparent stem reserves use during the effective kernel-filling period (ASRU_R2-R6_) for 14 cropping-system scenarios across 41 growing seasons (1970-1971 to 2011-2012) in Pergamino, Argentina. Data for estimation of ASRUR2-R6 were obtained from simulations performed with the Decision Support System for Agrotechnology Transfer (DSSAT)-CERES-Maize model. The analysis encompasses seven initial plant available soil water (PASW) × nitrogen (N) scenarios for early (15-Sep) and late (15-Dec) sowing dates: (i) potential (no water or N restriction), (ii) rainfed, with the whole profile (180 cm) at field capacity or 50% of PASW at sowing without N restriction (Full N), and (iii) the same PASW conditions as in (ii) but without N fertilization during the cycle and with initial soil N levels in the uppermost 45 cm as established by soil analyses in our experiments (i.e. 55 (N55) or 97 (N97) kg N ha^-1^ in early sowings and 80 (N80) or 142 (N142) kg N ha^-1^ in late sowings). The bolded numbers within the horizontal-colored bars indicated the percentage of cases when the R2-R6 stem biomass loss was (i) higher than the estimated ASRU_R2-R6_ (red bars), (ii) lower than the estimated ASRU_R2-R6_ (yellow bars), or (iii) null, as well as ASRU_R2-R6_ (green bars).

As compared to early sowings under potential conditions (i.e. when water or N were not limiting growth), delayed planting to mid-December (i) decreased the proportion of cases with null ASRU_R2-R6_ to only 17%, and (ii) increased to 34% the proportion of cases with estimated stem biomass loss during active kernel filling larger than the estimated levels of WSCS at R2 (Fig. 7). Moreover, lack of N fertilization did not modify the proportion of cases with null reserves use (20%-22%) or total reserves depletion (19-22%) for crops under no water deficit in late plantings. By contrast with early sowings under dryland farming, the model estimated that in late sowings (i) the proportion of cases with stem biomass drop > ASRU_R2-R6_ should be larger under full N (34%) than under no added N (22-27%), with almost no differences caused by the evaluated variations in initial soil N level (N80 and N142) and/or initial PASW (50% or 100%), and (ii) the proportion of cases with null ASRU_R2-R6_ should be larger under no added N (22%) than under full N (15-17%).

## 4. DISCUSSION

The present research integrated a set of variables that modulate KW determination for different end-use maize hybrids growing under contrasting environments. Together with the physiological determinants of KW (i.e. rate and duration of kernel filling), we analyzed the source/sink ratios and the water-soluble carbohydrates in stem during the kernel-filling period. Despite recent changes in maize production in Argentina, where farmers broadly adopted late sowings promoting the expansion of maize crops in more marginal and diverse environments, research and breeding efforts have been focused on grain hybrids grown under early sowings, with less attention to dual-purpose and silage hybrids and late sowings (Otegui et al., 2021).

The contrasting environments generated by extreme sowing dates and N rates within a year (sowing date × N) allowed the detection of differences in KW and its physiological determinants. The N250 condition greatly increased KW under early sowings, with little effects under the late ones. The lower impact of N fertilization on KW under late sowings could be attributed to the high initial soil N availability (i.e., N-NO_3_^-^), attributable to the higher temperatures experienced by soils during the fallow period and the pre-silking stage for late than for early sowing dates, a condition that favors the mineralization of organic matter (Bruun et al., 2006; Caviglia et al., 2014; Coyos et al., 2018). Variations in KW were mainly explained by variations in the kernel-filling duration (*r* = 0.82; Fig. 5), attributable to a shorter extension of this period for N0 than for N250 conditions, a trend that was more pronounced for early than for late sowings (Fig. 3). The shortening of the kernel-filling period could be directly associated with N shortage, which has been identified as one of the primary factors contributing to maize yield gaps (Aramburu Merlos et al., 2015; Ciampitti and Vyn, 2014). On-farm research showed that N restrictions may limit the attainable yield by reducing KW, with almost negligible effect on the kernel number (Moises et al., 2024). Similar responses were found in current research, where N fertilization only increased the kernel number in one environment, but increased the KW across all the explored environments (Table 2). Previous research also suggested that N fertilization can directly contribute to the determination of potential KW (Paponov et al., 2020) due to (i) an increase in the number of endosperm cells during the lag phase and, consequently, in the final kernel weight at maturity (Olmedo Pico et al., 2019), and (ii) the improved N content of the vegetative-tissue, which is lower under N0 than N250 conditions and could explain most of the KW variation in experiments where different N-fertilization rates were tested (Olmedo Pico et al., 2023). In previous research conducted at a higher latitude environment of the humid Pampas region of Argentina, the source/sink ratio during the effective kernel-filling period decreased due to delayed sowing date and negatively affected final KW (Cirilo and Andrade, 1996; Bonelli et al., 2016). This source limitation is mainly explained by the pronounced drop in both temperature and irradiance during the effective kernel-filling of late sowings in these environments (Cirilo and Andrade, 1996; Tsimba et al., 2013). Such drop did not occur in the late-sowing environments of the current research (Table 1), in agreement with previous analysis for similar late sowing dates in the same region (Mercau and Otegui, 2015).

Hybrids were grouped as for grain, dual-purpose, or silage end-use based on ecophysiological criteria. Our hybrid classification partially differed from those of the respective seed companies, which commercialized the hybrid KW 4360 AS-GL Stack for silage instead of a dual-purpose. In the current research, the dual-purpose behavior was achieved by combining the highest KW among all hybrid types with the low kernel number characteristic of silage hybrids (Table 2). The high KW of dual-purpose hybrids was achieved through the combination of the highest maximum kernel water content (strongly and positively associated with KW, *r* = 0.89; Fig. 5) and the highest kernel-filling rates under early environments (also positively associated with the kernel weight, *r* = 0.45; Fig. 5; Table 2; Fig. S1), both indicators of high potential KW (Borrás et al., 2003). During the critical period, the source/sink ratio was higher for dual-purpose and silage hybrids than for the graniferous ones, due to higher biomass accumulated from V14 to R2 (13.4% across environments) and a lower KNP (12.0% across environments; Table 2). The opposite trend was observed during the effective kernel-filling period, when the source/sink ratio was higher for graniferous than for dual-purpose and silage hybrids due to higher biomass accumulated from R2 to R6 (54.7% across environments), which overcompensated the higher KNP presented for the former than for the latter. As a result, SSR_V14-R2_ and SSR_R2-R6_ were negatively associated (*r* = 0.50; Fig.5).

The water-soluble carbohydrates in stem (in %) declined during the effective kernel-filling period for all tested growing conditions and hybrids. In previous research (Uhart and Andrade, 1991), using graniferous hybrids grown at Balcarce, Argentina (37°45’S, 58°18W) at stand densities of 8.5-9.1 plants m^-2^ a severe decline in this trait was detected under a shading treatment. Similarly, in a recent comparison between an old and a new maize hybrid grown at a lower stand density (6.1-7.6 plants m^-2^) in Manhattan, KS, USA (39°08′ N, 96°37′ W), Fernandez et al. (2021) found no remarkable decline in WSCS during kernel filling, except for a greater WSCS remobilization of the old than the new hybrid late in this period. By contrast, Kruse et al. (2008) estimated a marked decline in soluble carbohydrates during kernel filling for all plant fractions of several cultivars used for silage under high latitude (53°18’N, 98°80’E) and high stand density (9.0-10.0 plants m^-2^) in Hohenschulen, Germany. This apparent controversy may be related to the particular genotype × management × environmental conditions evaluated in each case, as was demonstrated in current research by testing different management strategies across a historical series of climatic data. With this approach, we were able to estimate the relative relevance of sowing date, PASW and N availability on reserves remobilization for most frequent management practices and initial soil conditions in the environment under consideration, which is a critical aspect in the determination of plant standability for maize used for grain (Pendleton and Egli, 1969; Pinheiro et al., 1984) and plant quality for maize used for silage (Andrieu et al., 1993), as discussed next.

In the field experiments developed in the current research, the observed level of WSCS (in %) at the beginning of the effective kernel-filling period (R2) was significantly lower for late than for early sowings, but never reached 0% during the evaluated period (Table 3) as estimated for different simulated scenarios (Fig. 7). The lower percentage of WSCS at R2 for late sowings was not due to a reduced amount of WSCS in terms of biomass (g per plant; Table S3). Instead, it was attributable to a higher stem volume among late-sown plants (Table 3), which promoted a dilution in the WSCS when expressed in percentage. Remobilization of WSCS from R2 to R6 was higher for late than for early sowings, analogous to the strong reductions in the WSCS reported under artificial source reductions by defoliation or shading during the kernel-filling period (Barnett and Pearce, 1983; Jones and Simmons, 1983; Reed et al., 1988; Tollenaar and Daynard, 1982; Uhart and Andrade, 1991) or under high latitude environments (Kruse et al., 2008). Such conditions induced the remobilization of reserves from the stem to the growing sinks (kernels) generated by the unbalanced relationship between the demand for assimilates by the reproductive sinks and the crop photosynthetic rate (Uhart and Andrade, 1991). Water-soluble carbohydrates in stem across the effective kernel-filling period were slightly higher under non-fertilized respect to fertilized treatments, which could be attributed to (i) the lower KNP in N0 than in N250 conditions, yielding a lower sink demand, and (ii) a greater SSR_R2-R6_ for the N250 than the N0 condition, resulting in enough current photosynthesis to supply assimilates for the growing kernels for the former but not for the latter (Fernández et al., 2021; Table 2). The percentage of WSCS across the effective kernel-filling period did not differ greatly among hybrid types (Table 3). At the beginning of the effective kernel-filling period, WSCS tended to be slightly higher for silage hybrids, a trend that could be attributable to the presence in this group of the EXP 1517 bm3 RR2 hybrid, which is a *bmr* genotype. Higher amounts of glucose and pentose have been previously reported among BMR hybrids at this stage, which are the result of mutations in the lignin biosynthetic pathway (*brownmidrib 1* and *brownmidrib3*) that allow easy accessibility to these carbohydrates (Barrière et al., 1994).

Finally, for grain and silage purposes, it is necessary to take into account the predominance of reduced levels of WSCS at R6 under late sowings. For grain purposes, the reduced stem reserves turn stalks weak and prone to lodging (Pendleton and Egli, 1969), in a context where achieving harvest kernel moisture after physiological maturity usually takes ∼3 months (Chazarreta et al., 2023). This constraint represents a serious difficulty for the mechanical harvest and contributes to reduced final grain yield (Pinheiro et al., 1984) and quality. In particular, it increases the likelihood of Fusarium ear rot, which raises fumonisin levels, with negative effects on human and animal health documented globally (Dutton, 1996; Camargos et al., 2003; Abbas et al., 2006; Fandohan et al., 2006; Maiorano et al., 2009; Castañares et al., 2019). For silage purposes, less stem reserves could also be related to low silage quality (i.e. less digestibility) due to reductions in the relation between the WSCS and cell wall (Andrieu et al., 1993). Hence, farmers must take into account the root and the stem strength of hybrids to avoid agronomic difficulties associated with lodging and stem breakage (Bonelli et al., 2016) and also be careful about the stem quality of maize plants for silage purposes, particularly under late sowing environments which currently represent ∼50% of the total land cropped to maize in Argentina.

## 5. CONCLUSIONS

This research shed light on the physiological mechanisms underlying kernel weight determination and the dynamic of water-soluble carbohydrates in stem in maize hybrids with varying end-uses growing under contrasting environmental conditions. Nitrogen fertilization needs careful consideration as low fertilization rates may negatively impact final kernel weight and, consequently, grain yield, being these unfavorable responses greater for early than for late sowings. Among hybrid types, dual-purpose hybrids showed the highest kernel weight, attributed to their superior maximum kernel water content, which enabled this hybrid type to hold high kernel-filling rates across environments. A greater remobilization of water-soluble carbohydrates in stem was observed for late-sowing environments compared to the early-sowing ones. The reduced percentage of WSCS under late-sowing environments observed in this study has important implications at the farming level. For grain production, these stems with depleted reserves became fragile, increasing the susceptibility to plant lodging and therefore affecting grain yield and quality. Regarding silage purposes, low percentages of WSCS could negatively impact the silage feeding value. In this regard, crop growth simulations suggest that PASW may not be the main driver of expected differences in ASRU_R2-R6_ levels under dryland farming. Estimated trends in this trait seemed primarily driven by the contrasting growing conditions created by markedly different sowing dates, N fertilization, and initial soil N conditions.

## 6. SUPPLEMENTARY DATA

The following supplementary data are available at JXB online.

Table S1. Dates of main phenological events of the experiments carried out during this research.

Figure S1. Relationship between the maximum kernel water content with the lag phase duration, and the kernel-filling rate with the maximum kernel water content.

Table S2. Correlation coefficients for the 7 variables included in the principal component analysis.

Table S3. Probabilities associated with the correlation coefficients for the 7 variables included in the principal component analysis.

## Acknowledgments

The authors want to acknowledge Juan Ignacio Amas, Luis Blanco, Facundo Curin and Octavio Ghio Trebino for their valuable assistance with fieldwork. Santiago Alvarez Prado and Maria E. Otegui are members of the National Council for Research (CONICET). Yésica D. Chazarreta holds a CONICET graduatés scholarship.

## 7. Author Contributions

**Yésica D. Chazarreta:** Conceptualization, Investigation, Formal analysis, Writing-Original Draft. **Santiago Alvarez Prado:** Conceptualization, Supervision, Writing - Review & Editing. **Maria E. Otegui:** Conceptualization, Investigation, Writing - Review & editing, Supervision, Funding acquisition, Project administration.

## 10. Data availability

Data will be made available on request.

## Abbreviations

ASRU_R2-R6_: Apparent stem reserves use during the effective kernel- filling period
KNP: kernel number per plant
KW: kernel weight
PASW: plant available soil water
SSR_V14-R2_: source/sink ratio during the critical period (V14-R2)
SSR_R2-R6_: source/sink ratio during the effective kernel-filling period (R2-R6)
WSCS: water-soluble carbohydrates in stem.

